# Calmodulin acts as a chaperone during co-translational folding of the Kv7.2 channel Calcium Responsive Domain

**DOI:** 10.1101/2024.11.15.622191

**Authors:** Arantza Muguruza-Montero, Jack R. Tait, Sara M-Alicante, Ane Metola, Eider Nuñez, Janire Urrutia, Vanda Sunderlíková, Alexandros Katranidis, Gunnar von Heijne, Sander J. Tans, Alvaro Villarroel

## Abstract

*In vivo*, the majority of nascent protein chains must begin folding during translation in order to obtain their native structure. While the importance of co-translational folding has become increasingly clear, the specific mechanisms underlying the coordination between the ribosome, nascent chain and molecular chaperones are still uncertain. Here, we have constructed a model of the co-translational folding pathway of the calcium responsive domain (CRD) of the human neuronal K_V_7.2 human neuronal ion channel, showing that calmodulin (CaM) is crucial. By combining Force Profile Analysis and single-molecule force spectroscopy techniques, we found that CaM, in a calcium-dependent manner, affects early folding events involving three key α-helices in the CRD. In addition, this study suggests that CaM at early stages induces the formation of metastable hairpins, as a part of the co-translational folding pathway. These findings expand on the role of CaM as a key regulator of folding in eukaryotes: not only as an essential cellular signaling protein, but also as a *bona fide* co-translational molecular chaperone.

## Introduction

Protein folding is a notoriously complex problem. While certain proteins can spontaneously fold in solution, the majority deviate into non-functional misfolded structures in the absence of dedicated folding mediators known as molecular chaperones. These chaperones are indispensable for cellular viability^1^, facilitating *de novo* protein folding and maturation, translocation, complex assembly, disaggregation and subsequent refolding, as well as aiding in degradation. Recently it has become clear that many of the required protein folding processes *in vivo* already begin co-translationally, before synthesis of the protein is complete. Indeed, interactions with the tunnel and surface of the ribosome can serve to chaperone nascent polypeptide chains directly, as well as providing a hub for recruitment of other dedicated chaperones^2^.

Calmodulin (CaM) is a small and flexible protein consisting of two lobes (N- and C-lobe). CaM is the primary calcium (Ca^2+^) sensor in eukaryotic cells, transmitting Ca^2+^ signaling to a diverse array of targets which are unable to bind this ion directly. It is, therefore, the paramount Ca^2+^ sensor in eukaryotic cells, located ubiquitously across all cellular compartments^3^. In addition, CaM participates in a number of chaperone-like activities: it prevents protein aggregation when the C-termini of K_V_7.1-7.2 channels are produced in bacteria^4,5^; it is involved in protein subunit assembly, and in nuclear and endoplasmic reticulum (ER) translocation^4,6–10^; it maintains the translocation competence of small-protein precursors of metazoans, and limits improper interactions with other cytosolic polypeptide-binding proteins^10^. In particular, CaM binding to K_V_7 channels is required for correct channel trafficking to the plasma membrane^7,9,11–14^. In addition, mutations responsible for self-limited benign familial neonatal epilepsy (BFNE) and severe epileptic encephalopathy, located in the calcium responsive domain (CRD) of the K_V_7.2 channel, disrupt CaM binding, leading to diminished currents due to ER retention^12,15^, which may reflect its proposed role in folding of the CRD during translation^16^. Although CaM is known to be involved in these pathways, the exact mechanisms underlying its influence are largely unknown.

The CRD is located in the long cytosolic C-terminal region of the K_V_7.2 channel and is comprised of three α-helices: hA, hTW and hB. In complex with CaM, the CRD forms a soluble antiparallel hairpin, with the C-and N-lobes of CaM bound to hA and hB, respectively^4^. The relative movement of hA and hB is crucial for Ca^2+^ signaling^4,17^, while the function of hTW is unclear. NMR data suggest that this small helix is very dynamic^4^; indeed, the available K_V_7 structures have not revealed a direct interaction between CaM and hTW through the target CaM binding clefts (PDBs: 6FEG, 6FEH, 7CR3, 7CR4, 7CR7, 8J01, 8J02, 8J04, 8J05, 8W4U)^4,18,19^. In general, hTW is thought to be a backup option for K_V_7.2 function: whereas it is itself dispensable without disrupting channel function, it becomes essential when CaM docking sites at hA or hB are perturbed^20^.

The present work provides a comprehensive picture of the co-translational folding process of the CRD and highlights the critical role played by CaM in this intricate molecular ballet. In addition, our results suggest the transient formation of hairpins between hA and hTW and, subsequently, between hTW and hB, as part of the co-translational folding pathway in a calcium-dependent manner.

## Results

### Impact of calmodulin on the *in vivo* force profile of K_V_7.2 CRD

To pinpoint the key translation stages where folding is promoted by CaM, we employed Force Profile Analysis (FPA), taking advantage of the force-sensitivity of translational arrest peptides (APs) to monitor co-translational folding^21,22^. APs are sequences which bind in the interior of the ribosomal exit tunnel, preventing chain elongation. The stalling efficiency is highly dependent on the external force applied to the nascent chain when the final residue in the AP is translated^21,22^. Such pulling forces can be generated by folding of the nascent chain nearby, or inside, the ribosomal exit tunnel^21,22^. APs allow generating highly-sensitive biosensors for reporting on co-translational folding events: the fraction of full-length protein (f_FL_) serves as a proxy for the folding propensity of the newly-synthesised nascent chain at each stage of translation, as shown previously^23^ (Fig. 1A).

**Figure 1.**
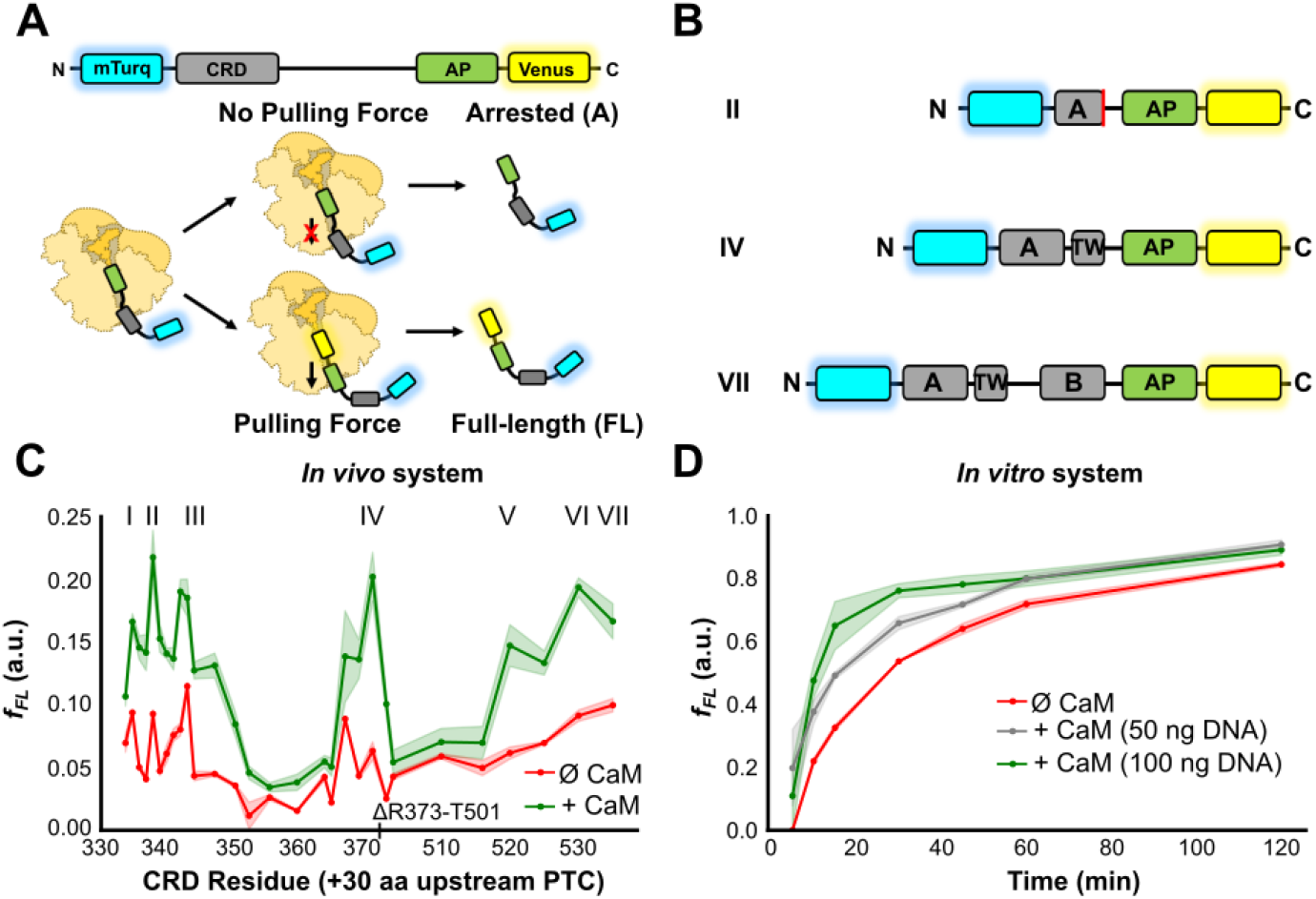
The *in vivo* force profile reveals the CaM dependency of three peaks, which correspond to the exit of the CRD helices from the ribosomal tunnel. **(A)** Schematic representation of the FPA using the fluorescent variant on top. Blue and yellow fluorescent proteins have a halo indicating its fluorescence. CRD: Calcium Responsive Domain; AP: Arrest Peptide. **(B)** Schematic representation of variants associated with the indicated peaks in the *in vivo* FPA. The red line on the scheme of the II variant indicates that hA is incomplete. **(C)** *In vivo* force profiles for the CRD alone (red) and after co-expression with CaM (green). A schematic representation of the corresponding CRD is shown above. Shaded regions represent SEM (n = 4-12). **(D)** *In vitro* time-course pulling-force assay of the CRD without CaM (red) and with varying amounts of CaM cDNA (50 ng in gray and 100 ng in green). Shaded regions represent SEM (n = 3).

Here, we generated a library of 29 variants containing K_V_7.2 CRD sequences of increasing length, flanked by an N-terminal mTurquoise2.1 fluorescent protein, and a C-terminal mcpVenus173 fluorescent protein downstream of the SecM (*Ec-Ms*) AP^24^ (Fig. 1B) (from now on referred to as Turquoise and Venus, respectively). In this experimental setup, two main products are obtained: an arrested (A) product from the protein labeled only with Turquoise, and a full-length (FL) protein that is labeled with both Turquoise and Venus (Fig. S4). This enabled measurement of f_FL_ by the ratio of yellow:blue fluorescence intensity, which in turn allowed us to directly monitor the influence of CaM on folding of nascent K_V_7.2 CRD proteins during translation.

*In vivo* co-expression of the library constructs with CaM resulted in a force profile with three clear folding transitions (marked by increases in f_FL_ at peaks I-III, IV and V-VII; Fig. 1C, green). Considering the length of the ribosome exit tunnel (≈30 residues^25,26^), these transitions correspond to the sequential exit from the tunnel vestibule of each of the three α-helices comprising the CRD. In particular, the first transition (corresponding to folding of helix hA) is split into three sub-peaks (I, II and III). Notably, sub-peaks II and III align with the emergence of two hydrophobic residues that are pivotal for CaM binding within the IQ motif of hA (IQSA<u>W</u>R, hydrophobic residues are underlined) in K_V_7.2 (Fig. S1)^27^. Crucially, in the absence of CaM, we observed a global decrease in folding force across the entire CRD (Fig. 1C, red). This implies that while the three α-helices retain some capacity to fold independently, these transitions are significantly dependent on the presence of CaM.

To explore folding in the absence of cellular context, *in vitro* FPA was performed. Similarly to *in vivo, in vitro* FPA showed an f_FL_ increase at almost every translation stage in the presence of CaM (Fig S4). Both profiles were qualitatively similar, in terms of CaM dependency and number of major peaks, although the resolution for the *in vivo* FPA was more favorable.

We addressed how CaM abundance affected the process *in vitro* (Fig. 1D), using a variant which ensured the complete exit of the whole CRD (L21, see materials and methods, Fig. S1). After 2 hours, we observed accumulation of FL protein even in the absence of CaM expression (Fig. 1D, red line). Significantly, the production of FL protein accelerated in a CaM-dependent manner (Fig. 1D). These results reinforce the concept of a role of CaM in assisting CRD co-translational folding and suggest the existence of three steps, corresponding with the sequential exit of hA, hTW and hB from the ribosomal tunnel.

### Ca^2+^ binding to CaM impacts on co-translational folding

The role of CaM as a crucial Ca^2+^ modulator in various physiological pathways prompted us to investigate the role of Ca^2+^ binding. For this purpose, we employed two CaM mutants unable to bind Ca^2+^ at the N-lobe (CaM12) or the C-lobe (CaM34). We used a reporter with a 21-residues-long tether, previously found to correspond with a CaM-dependent peak in the FPA (from now on referred to as the CRD L21 co-translational folding biosensor^16^). The results revealed a significant signal reduction of the full-length fraction for both CaM12 and CaM34 versus WT CaM (Fig 2A). However, folding was not totally abolished: the fraction of full-length protein remained significantly higher in both mutants than in the absence of CaM, suggesting that while the ability of CaM to bind Ca^2+^ is beneficial in enabling folding of the CRD, it is not strictly required.

**Figure 2.**
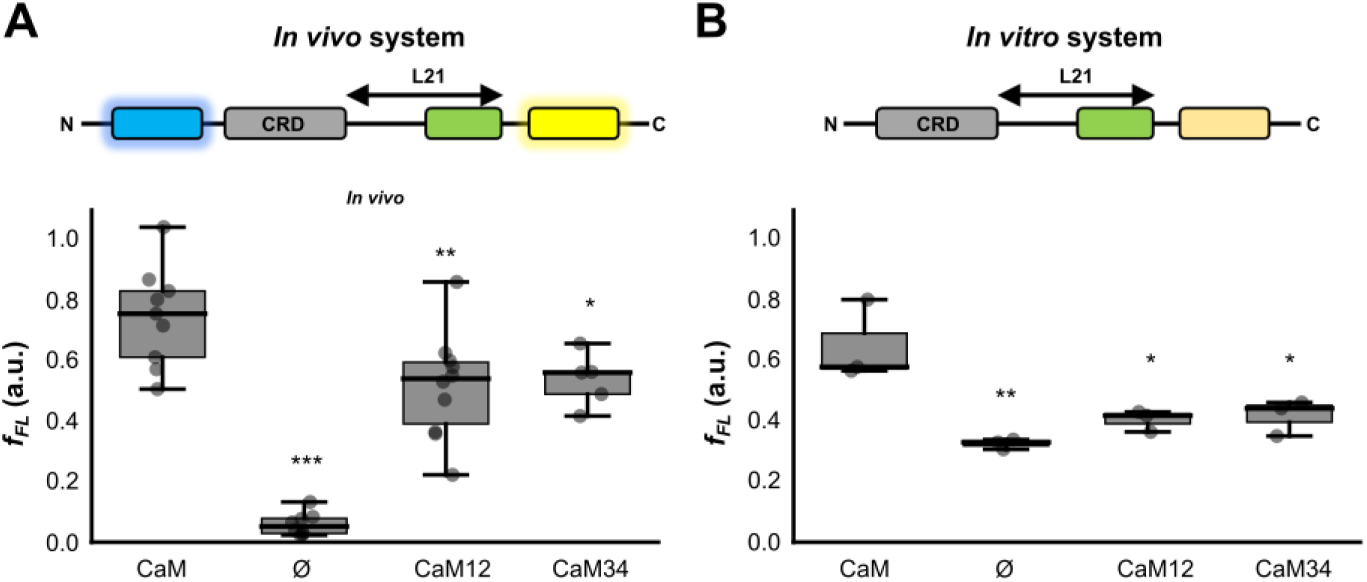
*In vivo* and *in vitro* pulling-force assays with CaM variants unable to bind Ca^2+^ at the N- or C-lobes. **(A)** Box plot indicating the f_FL_ of the CRD L21 folding biosensor *in vivo* without CaM and co-expressed with CaM, CaM12 and CaM34. Whiskers indicate SEM (n_CaM WT_ = 9; n_ø_ = 8; n_CaM12_ = 10; n_CaM34_ = 5). **(B)** Box plot indicating the f_FL_ of *in vitro* assay of the CRD L21 folding biosensor alone and co-expressed with CaM lobe mutants at 1:1 ratio. Whiskers indicate SEM (n = 3). A schematic representation of the variant used in each system is shown above. (***p ≤ 0.001, **p ≤ 0.01, *p ≤ 0.05 vs. CaM).

*In vitro* similar results were obtained: the folding force of the CRD was decreased, but not completely abolished, for both CaM12 and CaM34 (Fig. 2B).

### hA has a preponderant role in folding of the CRD

After establishing the overall impact of CaM on CRD folding, we aimed to elucidate the native pathway followed by the nascent CRD during translation. We focused on two key reporters corresponding to peaks IV and VII (Fig 1), associated with exit of hTW and hB from the ribosomal tunnel, respectively. For each construct, each α-helical domain was disrupted with GSG stretches, to prevent folding, as indicated (Fig. 3). At stage IV, where hTW presumably is just outside the tunnel, we found that CaM-mediated folding of hTW is strongly dependent on proper formation of hA: GSG disruption of hA caused a significant reduction in the full-length protein production, even though CaM was also present (Fig. 3A). Disruption of hTW itself also caused a comparable reduction in f_FL_. The folding transition observed at stage IV must therefore involve folding of both helices (in complex with CaM). At stage VII, we found that folding of hB is in turn highly dependent on proper formation of hTW: GSG disruption of hTW resulted in f_FL_ values indistinguishable from those obtained for the WT in the absence of CaM (Fig. 3B). Given that we already established that hTW is itself dependent on hA, one would expect that hB folding is also (indirectly) dependent on hA. Indeed, f_FL_ measured on the hA-disrupted VII construct was partially decreased.

**Figure 3.**
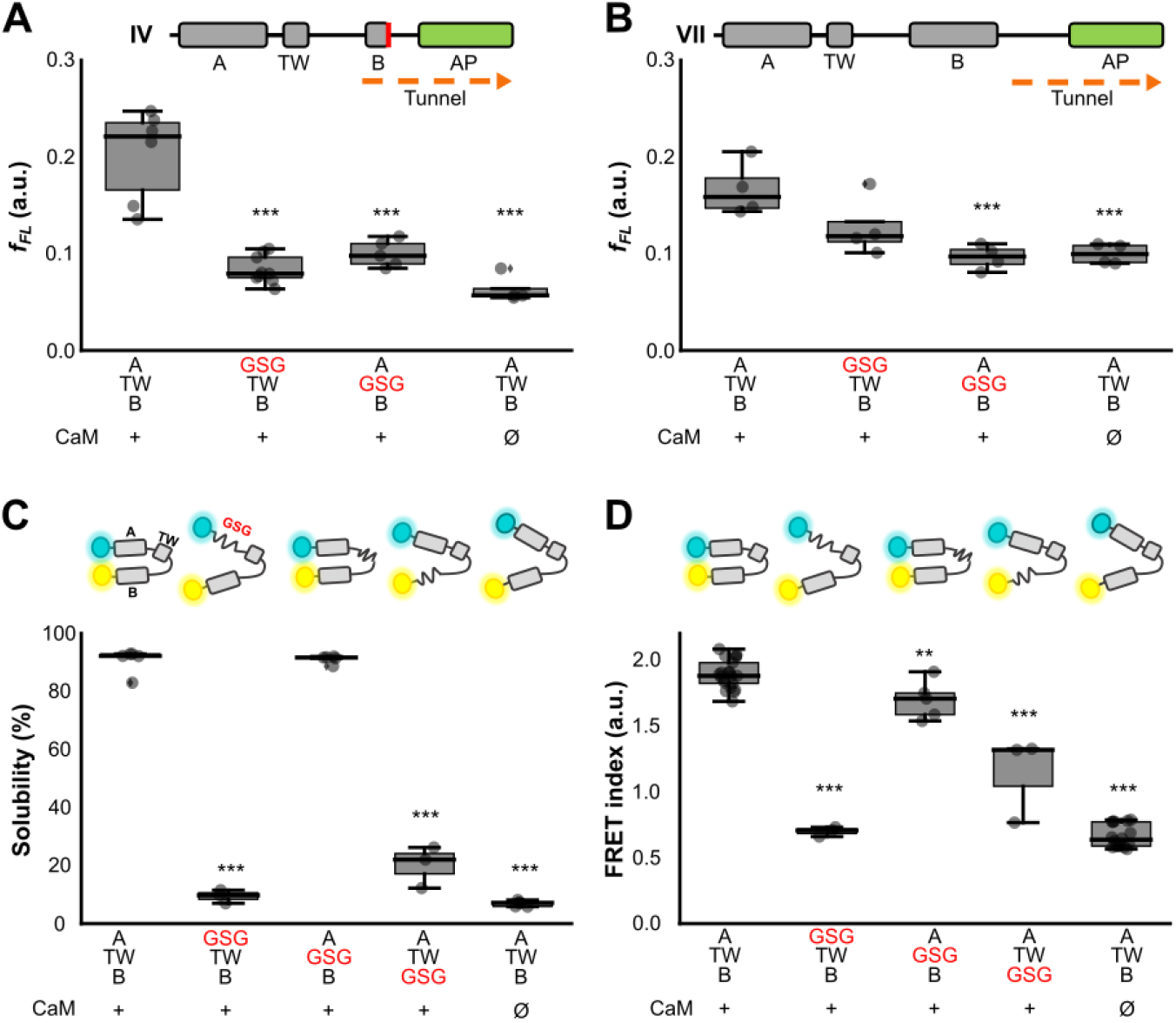
Importance of each of the CRD helices folding. **(A)** Box plot indicating the f_FL_ of the variant corresponding to peaks IV and **(B)** VII with a GSG linker substitution in hA or in TW co-expressed with CaM. (n = 4, except for n_IV(WT+CaM)_ = 6, n_IV(GSG in hA)_ = 9, n_IV(GSG in hTW)_ = 5). Schematic representation of the WT (A-TW-B) variant corresponding to IV and VII is indicated at the top. **(C)** Solubility: Box plot of the percentage of the fluorescence intensity of the supernatant band of SDS-PAGE gels. (n = 3, n_GSG_ for hTW = 5). **(D)** Box plot of the FRET index: Values obtained from spectra of the soluble fraction. (n = 3, 5, 3, respectively for hA-, hTW- and hB-disrupted-CRD). In the upper part of both panels, A and B, there is a schematic representation of the biosensors. (n = 3, except for n_GSG in hTW_ = 5). Asterisks refers to the significance compared to the WT variants (***p ≤ 0.001 and **p ≤ 0.01) for all the pannels.

The native CRD is known to form an antiparallel hairpin in the presence of CaM^4^. By measuring protein solubility and Förster Resonance Energy Transfer (FRET) between the N- and C-terminal fluorophores (Fig. 3C and D, respectively), we can thereby follow the formation of such antiparallel hairpins *in vivo*. We used this concept to monitor structure formation of the CRD upon perturbation of each α-helix (Fig. 3C and D). Disruption of the small hTW had little effect on the native hairpin: FRET was only marginally decreased, and solubility remained unchanged. This is consistent with existing structural models, where hTW does not make close contacts with CaM^4,28^. As expected, no hairpin formation was observed when hA was perturbed - both FRET and solubility were completely abolished. However, the same was not true for hB perturbation: although both FRET and solubility measurements were significantly reduced, they remained above the baseline reference in absence of CaM. This suggests the formation of a metastable hairpin between hA and hTW in the presence of CaM: the partial fold would allow moderate solubility, while also ensuring a 3D conformation that confers some energy transfer.

### CaM binding induces novel hairpin formations

Next, we conducted an optical tweezers force spectroscopy assay to probe the conformation of single, nascent CRD molecules at nanometer resolution, both in the presence and absence of CaM (Fig. 4A). We generated stalled ribosomes expressing CRD variants using *in vitro* transcription-translation^29^. Polystyrene beads were functionalized with stalled ribosomes using DNA “handle” linkers. By immobilizing each bead in a steerable optical trap, we could drive the inter-bead distance to repetitively stretch (Fig. 4B, light green) and relax (Fig. 4B, dark green) the nascent chain. Unfolding of the nascent chain during this stretching was marked by sharp discontinuities in the resulting force-extension curve, characterized by specific contour lengths (L_C_) and unfolding forces (Fig. 4B, grey arrows). This mechanical control allowed us to probe the conformation space of the nascent CRD in the presence and absence of CaM, as a function of both nascent chain length (i.e. IV and VII translation stages). Single-molecule variants of these constructs (IV_SM_ and VII_SM_ respectively) were generated by inclusion of an amber codon and GSG linker, to enable tethering of the nascent chain.

**Figure 4.**
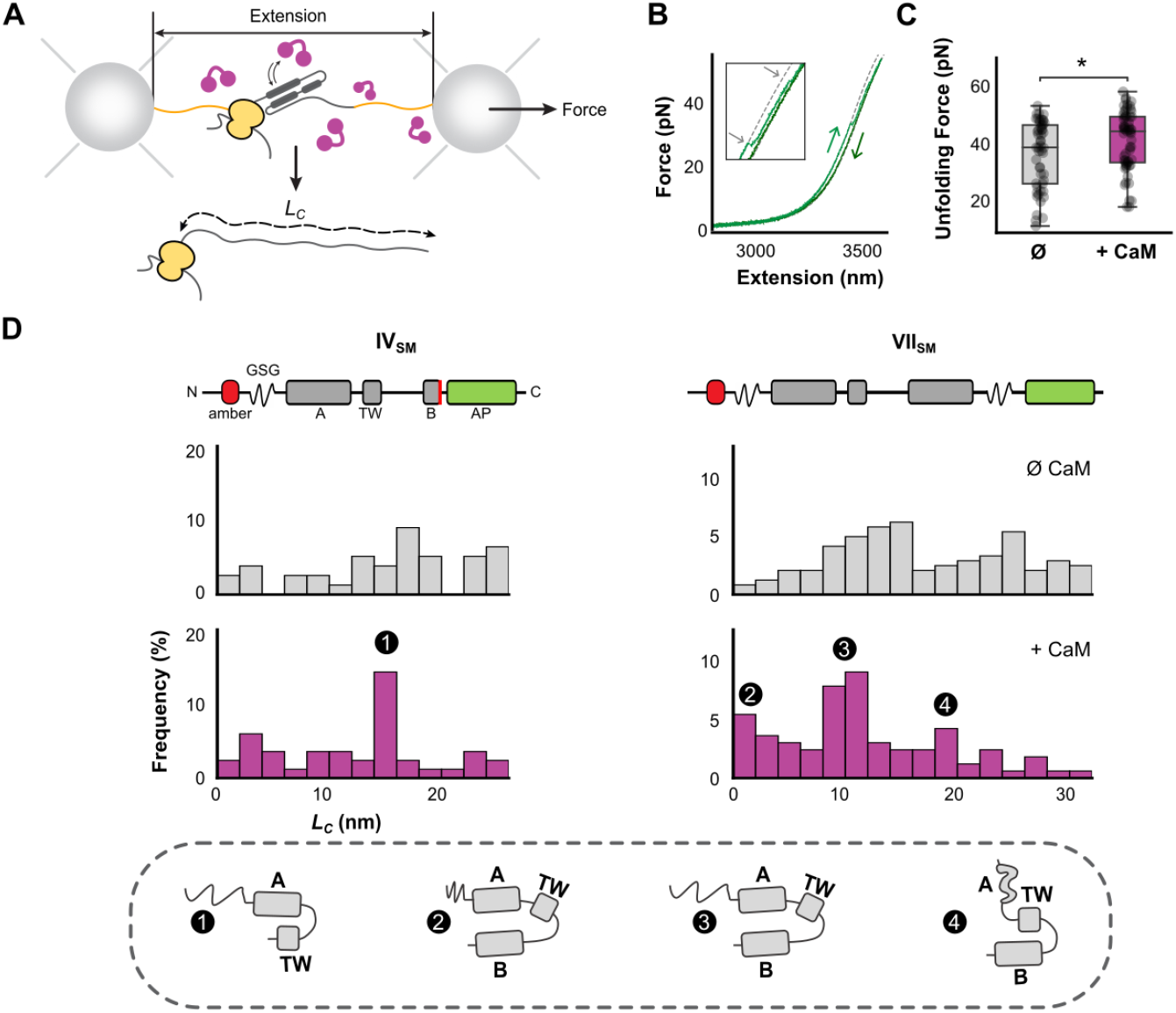
Identification of three CaM-dependent unfolding states of nascent CRD. **(A)** Schematic representation of the experimental setup, showing a ribosome expressing the CRD (green) tethered between two optically-trapped beads using DNA handles (orange). These experiments were performed both with and without added CaM (purple). **(B)** Example force-extension data showing unfolding of the nascent CRD VII_SM_. In the experiment, the nascent chain is repetitively pulled (light green curve) and relaxed (dark green curve). From these force-extension data, unfolding events can be identified (inset, grey arrows), and the unfolding force (F) and the end-to-end length of the unfolded part of the protein (L_C_) can be quantified. **(C)** Forces for all unfolding events of CRD VII_SM_ both with and without CaM. Binding of CaM to the nascent CRD stabilizes the domain against mechanical unfolding. **(D)** Histograms showing all observed folding states, based on the Lc values measured from the force-extension data, in the absence and presence of CaM (grey and purple, respectively) for the IV_SM_ and VII_SM_ variants (schematic representations of the two constructs are presented at top). Peaks are present at 14.9 nm for the IV_SM_ construct; and at 0, 10.2 and 17.7 nm for the VII_SM_ construct (Figure S5, Table S1). Bottom, schematic representation of the possible structures identified and described. States identified by force spectroscopy are consistent with the following structures: 1, a hairpin formed between hA and hTW (IV_SM_); 2, a fully compacted state with the entire CBD folded (VII_SM_); 3, the “native” state (VII_SM_) with the entire CRD folded and the N-terminal tail fully extended; 4, a hairpin formed between hTW and hB, where the hA helix is unfolded and fully extended (VII_SM_). N-values detailing number of experiments are given in Methods.

Efficient binding of CaM to the nascent CRD in the single-molecule assay was confirmed by an overall increase in the average domain unfolding force (Fig. 4C), consistent with mechanical stabilization of the nascent domain due to interactions with CaM. In the absence of CaM, the conformation of the nascent CRD was highly variable, regardless of translation stage: at both stage IV and VII we observed a relatively uniform ensemble of states, ranging from fully-compacted (L_C_ = 0) to fully-unfolded (L_C_ = 28 / 34 nm; Fig. 4D). However, when CaM was added, clear states emerged. At stage VII, CaM specifically promoted two states, with L_C_ ≈ 1 nm (or negligible) and L_C_ ≈ 10 nm respectively (Fig. 4D, right). Using the known structure of the CaM/CRD complex^4^, we could compute the expected contour lengths of various CRD conformations (see Methods). In this case, both emerging states are consistent with the native hA-hB antiparallel hairpin conformation: the 1 nm state maps to the crystallographic distance between the termini of the α-helices; while the 10 nm state is consistent with the same core hairpin, but where the intrinsically-disordered N-terminal domain^4^ is extended by the applied force (Fig. 4D, bottom). In addition, there was also a less-abundant state at 18 nm which perfectly maps to a hTW-hB hairpin.

At stage IV, although the L_C_ distribution in the absence of CaM is as homogeneous as in stage VII, one clear state emerges in the presence of CaM with L_C_ ≈ 15 nm (Fig. 4D, left). This perfectly matches the expected contour length of the proposed hA-hTW antiparallel hairpin (after accounting for differences in the extended length of the unstructured C-terminal region between the two constructs). Notably, this same state in the VII distribution (Fig. 4D, left column 1; L_C_ ≈ 25 nm, due to C-terminal length differences) is diminished, supporting the earlier observation that the presence of hB prevents CaM forming the hA-hTW intermediate.

To obtain further insights, we modified the WT FRET reporter by eliminating hB or hA, leaving the hA-hTW or hTW-hB sequences flanked directly by the two fluorophores, respectively. Expression of the hA-hTW construct in presence of CaM resulted in a FRET index indistinguishable from that of the WT reporter (Fig. 5B), with only moderately-decreased solubility (Fig. 5A), again indicating that CaM can indeed stabilize an antiparallel hairpin between hA and hTW alone. However, FRET and solubility were significantly reduced for the hTW-hB construct in the presence of CaM, probably indicating that this hairpin is destabilized once the final protein product is synthesized and released from the ribosome (Fig. 5).

**Figure 5.**
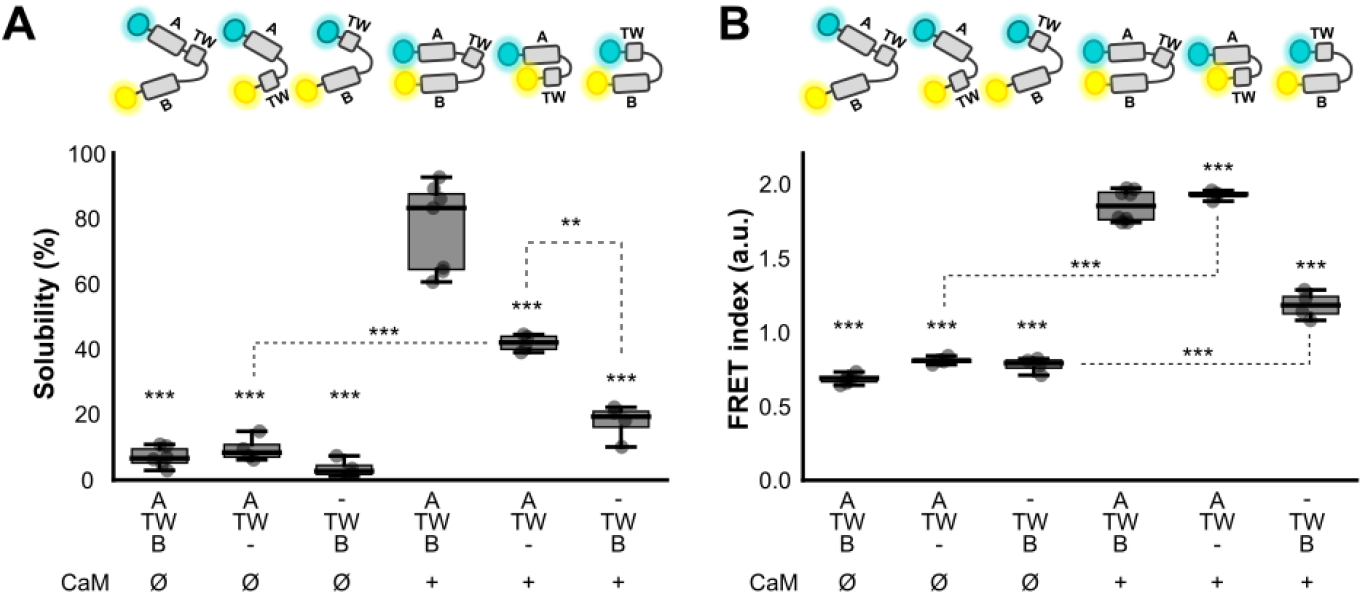
hTW forms a hairpin with hA in the presence of CaM. **(A)** Solubility: Fluorescence intensity of the supernatant band in SDS-PAGE gels for the hA-hTW-hB, hA-hTW and hTW-hB biosensors without and with co-expression of CaM. (n=4, n_hA-hTW-hB+CaM_=7, n_hA-hTW-hB_=6). **(B)** FRET index: Values obtained from spectra of the soluble fraction. (n=4, n_hA-hTW-hB+CaM_=8, n_hA-hTW-hB_=6). Asterisks located at the top of each box refer to the significance compared to the WT biosensor in the presence of CaM (***p ≤ 0.001 and **p ≤ 0.01).

In addition, we employed AlphaFold Multimer to predict the three-dimensional conformations of these nascent chains at each arrested position^30,31^. The predicted structures were analyzed and selected based on consistency with our experimental data, revealing a plausible series of folding events for the nascent K_V_7 domain.

Figure 6 provides a schematic representation of this proposed co-translational folding pathway. In this model, the prefolded state of helix A (hA) is shown in orange, helix TW (hTW) and helix B (hB) are colored in green and blue, respectively, while calmodulin (CaM) molecules are depicted in magenta. The middle states along the folding pathway were derived from AlphaFold Multimer predictions, corresponding to the sequences tested in our assays. These intermediates highlight critical stages of nascent chain folding, revealing the progressive structuring of the helices and the engagement of CaM as translation proceeds.

**Figure 6.**
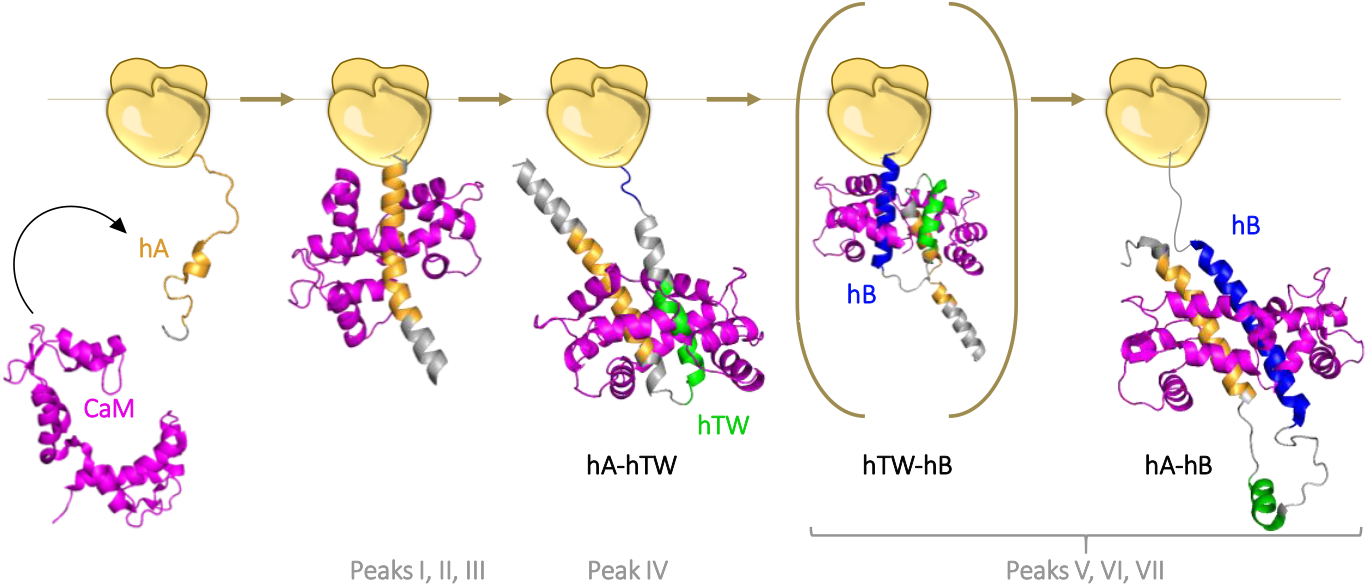
Schematic representation of a plausible co-translational folding pathway. hA, hTW and hB are colored in orange, green and blue, respectively. CaM molecules are colored in magenta. The prefolded state of hA (left) was based on the prefolded state of the IQ-like motif of SK2, since the structure of hA-K_V_7.2 in the absence of CaM is not available (PDB: 1KKD). The middle states were obtained from running AlphaFold Multimer predictions of different sequences used for FPA and CaM. The final state (right) was obtained from the NMR structure of the K_V_7.2 CRD/CaM complex (PDB: 6FEH).

The final folded state illustrates the culmination of co-translational folding, in which CaM is tightly bound to the fully formed helices, stabilizing the mature conformation of the K_V_7 domain. Overall, these data suggest a stepwise, co-translational folding process, beginning with the initial formation of hA and culminating in the complete assembly of the K_V_7.2-CaM complex.

## Discussion

In this study, we perform a comprehensive analysis of the role of CaM in mediating the co-translational folding of the K_V_7.2 CRD. Firstly, we show that CaM enhances the stability of the nascent CRD during translation, as evidenced by increased folding propensity (Force Profile Analysis, Fig. 1C), increased release rate of the arrested state during translation (time-course force-pulling assay, Fig. 1D), and increased mechanical stability (force spectroscopy assay, Fig. 4C). This stabilization may arise from protection of hydrophobic regions during protein synthesis, or by mitigating interactions with the ribosome that decelerate and destabilize nascent chain folding^32^. Secondly, interactions with CaM serve to alter the folding pathway of the substrate: in the absence of CaM the nascent chain forms a wide range of off-pathway intermediates, but with CaM bound it is reliably funneled towards the known functional hairpin conformation (Fig. 4D). In short, we demonstrate that CaM exhibits all the hallmarks of a *bona fide* molecular chaperone.

Given the role of CaM as a major regulator of Ca^2+^ signaling, we conducted experiments to address the role of this ion in co-translational folding. *In vivo*, Ca^2+^ signaling has already been proposed to regulate folding via a number of pathways, including the activity of the ribosome^33–36^ and chaperones in the ER^11,12^. Here, we suggest another layer of complexity, whereby the chaperone activity of CaM may itself be regulated by Ca^2+^ signaling. Indeed, we demonstrate here that the ability of CaM to bind Ca^2+^ is important for its chaperone function (Fig. 2), which suggests the existence of an important interplay between signaling and chaperoning in general. The exact role of Ca^2+^/CaM in folding is an area that warrants further investigation.

There is substantial evidence that Ca^2+^ signaling through CaM regulates the exit of K_V_7.2 from the ER^11,12^. While CaM binding is a prerequisite for ER exit, binding by itself is not sufficient for this to happen^11^. Since proteins that are not properly folded fail to exit the ER, it follows that ER exit regulation by CaM may be a consequence of its role in co-translational folding. Interestingly, it has been shown that CaM can modulate the Sec61-mediated translocation pathway^10,37^. It is therefore reasonable to propose that CaM might also be involved in the modulation the translocation of K_V_7 across ER membrane during the synthesis through a similar mechanism.

This study helps to clarify the elusive role of hTW. While classical binding experiments have shown convincingly that hA and hB peptides bind CaM, we have failed to detect any similar interaction between CaM and hTW peptide (data not shown). Here, we demonstrate that this helix has an impact in CaM-dependent folding (Fig. 1C). The formation of CaM-dependent hA-hTW and hTW-hB hairpins detected in single-molecule pulling experiments suggest a direct interaction between CaM and hTW (Fig. 4D), which is, indeed, predicted by AlphaFold (Fig. 6). Regarding the functional role of hTW, removal of this helix has minimal impact on the electrophysiological properties of K_V_7.2 channels^20^. However, hTW is essential for function when its binding to either hA or hB is weakened by specific mutations. Consistent with the functional data, our results reveal that hTW is not essential for hA-hB hairpin formation. Indeed here, there was little or no effect on solubility or FRET after hTW disruption (Fig. 3C-D). Nevertheless, we demonstrate that it is crucial for mediating co-translational folding: formation of hA was necessary for hTW stability (and *vice-versa*); formation of hTW was required for hB stability (Fig. 3C); and force spectroscopy and FRET experiments revealed a number of intermediate folding states all involving hTW (Fig. 4, Fig. 5). In other words, disruption of hTW appears to reduce the probability of following the proper folding pathway towards a “productive” conformation, but does not totally impede formation of the native fold. This may explain why mutations at hTW are associated with human disease and alter CaM binding affinity^20^; and why K_V_7.2 channels lacking hTW exhibit a more heterogeneous surface expression profile^20^. Furthermore, binding studies have demonstrated that the CRD binds more favorably to CaM than an equimolar mixture of individual hA and hB peptides^38^, reinforcing the idea that hTW may stabilize the complex.

Through the fruitful combination of force profile analysis, FRET measurements, single-molecule force spectroscopy, and previous molecular dynamics simulations^27^, we have assembled evidence to suggest a plausible mechanism for the co-translational folding of the K_V_7.2 CRD (Fig. 6). First, CaM is recruited to the nascent CRD through recognition of a pre-helical “capping” conformation of hA (possibly shaped by interactions with the ribosomal tunnel^39,40^, as previously reported^27,41^) (peaks I-III, Fig. 1 C). As hTW exits the ribosome, it forms a transient hairpin intermediate with hA that is embraced by CaM (peak IV, Fig. 1 C), stabilizing the nascent chain. Finally, as hB begins to emerge, the hA-hTW hairpin is destabilized and hB replaces hTW in the hairpin (peaks V-VII, Fig. 1 C), completing the native fold. Prior to the final hairpin stabilization, a hairpin between hTW and hB could be formed (peaks V-VI, Fig. 1 C).

To provide more support for this model we used AlphaFold Multimer to predict likely structures. Interestingly, the hA-hTW and hTW-hB hairpins suggested by our results were anticipated in complex with CaM (Fig. 6)^30,31^. Once hB exits from the ribosomal tunnel, as indicated by single molecule unfolding (Fig. 4) and helix dependency results (Fig. 3C), hTW has the option to form a hairpin with either hA or hB. According to this model, hB would subsequently replace hTW in the final hA/hB hairpin, which is in turn stabilized by CaM (Fig. 6).

We demonstrate CaM chaperone activity for the K_V_7.2 CRD folding. The principles we propose could be generalizable to many protein classes across the eukaryotic proteome, particularly given the broad range of natural targets of CaM *in vivo*. Future research is bound to reveal which other proteins or domains are chaperoned by CaM during their synthesis, and to underscore the implications in folding-related diseases.

## Materials and Methods

### Library of co-translational folding biosensors

To perform the Force Profile Analysis *in vivo*, twenty-nine constructs were created by sequentially shortening the CRD residue by residue at regions of interest (Fig S1). CRD domains of variable length were followed by the arrest peptide (AP) SecM (*Ec-Ms*) (FSTPVWISQHAPIRGSP) and flanked by mTurquoise2.1 and mcpVenus in the N- and C-terminal, respectively. These constructs were cloned into the pProEx-HTc by GenScript. Helices A or TW from CRD were then replaced by a GSG linker in some of these constructs (corresponding to peaks IV and VII from the FPA, Fig 4) (Fig S2).

These variants were cloned into the pET19b plasmid using Gibson Assembly for *in vitro* transcription/translation. Since the N- and C-terminal of the designed constructs were identical, this technique enabled high throughput cloning using the same four primers for all inserts (see Supplemental text 1 for further information about Gibson Assembly). In this clones the mTurquoise2.1 was removed, starting directly from the CRD, and the C-terminal mcpVenus was replaced by a LepB P2 domain-derived sequence encoding 23 residues (GSSDKQEGEWPTGLRLSRIGGIH), to enable radioactive quantification.

To analyze the fraction of full-length ensuring that the entire CRD has exited from the ribosomal tunnel, an already described co-translational folding biosensor was used^16^. This consists of the CRD followed by a 21-residue-long linker including the 17 residues of the SecM (*Ec-Ms*) (EFYV-FSTPVWISQHAPIRGSP) (referred to as L21) coding sequence and flanked by mTFP1 and mcpVenus fluorescent proteins in the N- and C-termini, respectively, and cloned into pProEX-HTc plasmid (Fig S1). This variant was also cloned in pET19b plasmid for *in vitro* transcription/translation.

Note that all these co-translational folding biosensors were used to calculate the fraction of full-length which reports on folding during translation as the fraction of full-length. However, to analyze the final folding of the protein other biosensors were used (folding biosensors) using solubility and FRET index as measurements. The WT folding biosensor, which was already described^5^, consists on the CRD flanked by mTFP1 and mcpVenus fluorescent proteins in the N- and C-termini, respectively, and is cloned into the pProEX-HTc plasmid. This biosensor reports on the folding of the hairpin between hA and hB is indicated as an increase in FRET index. In addition, each helix of the CRD was substituted by a GSG linker in independent folding biosensors (Fig S3).

### Mutant calmodulin versions

CaM12, CaM34, and CaM1234 in pOKD4 vector were synthesized by GenScript Biotech Corporation (Netherlands). CaM12 has mutation in the EF1 and EF2 (D22A and D58A, respectively), CaM34 in the EF3 and EF4 (D95A and D131A, respectively) and CaM1234 has mutations in all the EF-hands (D22A, D58A, D95A, and D131A, respectively).

### *In vivo* expression for Force Profile Analysis (FPA)

The library of pProEX-HTc plasmids coding increasing lengths of K_V_7.2 CRD constructs was transformed in *E. coli* BL21 DE3 cells, either alone or with the pOKD4 plasmid carrying the CaM coding gene. Cells were grown overnight at 37 °C from single colonies, and then diluted into 10 ml of fresh LB (1:100 dilution) and incubated at 37 °C until reaching an OD_600_ of 0.6. Protein expression was induced during 3 h at 37 °C by the addition of 0.5 mM IPTG. Cells were harvested by centrifugation at 5478 g (7000 rpm with Eppendorf F-35-6-30 Rotor for 5430/5430R rotor) for 5 min. The cell pellets were resuspended in 500 µl lysis buffer (50 mM HEPES, pH 7.4, 120 mM KCl, 5 mM NaCl, 5 mM EGTA, 0.5 mM dithiothreitol (DTT), and protease inhibitors (1X Complete; Roche Applied Science), and 1 mM PMSF), and similar OD values were fitted for all the samples. The cellular cultures were sonicated using 3 cycles of 10 s on / 10 s off, and centrifuged at 19,000×g during 30 min at 4 °C for supernatant and pellet separation. The pellets were resuspended in the same buffer volume used before.

The soluble protein fractions were analyzed in a Fluoromax-3 fluorometer by recording the emission spectra of mTurquoise or mTFP1 and mcpVenus fluorescent proteins upon excitation at 440 or 458 and 515 nm, respectively.

In this experimental setup, the synthesis outcome of these variants or folding biosensors is indicative of the pulling force exerted on the nascent chain and related to the folding status during translation. If the protein is stalled, an arrested protein (A) will be produced. In contrast, if folding of the partial CRD occurs during translation the full-length protein (FL) will be synthesized (Fig 1A). Thus, quantifying the stalling efficiency (f_FL_) provides a measure of co-translational folding events. This is computed as the ratio between the emissions at the peak wavelength for Venus and for mTurq or mTFP1. This analysis is conducted after a 3-hour expression period, both in the presence and absence of CaM, within *E. coli*.

The fraction of full-length (f_FL_) in the FPA was studied by SDS-PAGE electrophoresis (15% acrylamide gels) using unboiled samples. The gels were visualized using Versadoc imaging equipment, exciting using blue or green LEDs, combined with 530BP28 or 605BP35 emission filters. After merging both images, the in-gel f_FL_ was computed as the proportion of FL present in the soluble samples (f_FL_ = I_FL_ = I_FL_/(I_FL_+I_A_)), by quantifying the fluorescent intensity of the bands corresponding to full-length (I_FL_) and (I_A_) using FIJI (ImageJ) software.

### *In vitro* expression for FPA

*n vitro* expression and analysis were conducted as previously described^42^. Briefly, a linear DNA product was generated from each construct plasmid by PCR using Q5 polymerase with forward and reverse primers that anneal to the T7 promoter and terminator regions, respectively. Following PCR cleanup (using the manufacturer’s instructions), the product was confirmed by agarose gel electrophoresis. *In vitro* transcription and translation were carried out in the PURExpress commercial system (mixed according to the manufacturer’s recommendations). One hundred nanograms of the PCR product of K_V_7.2 CRD constructions were added into the reaction and *in vitro* co-expression of CaM was carried out in a CRD:CaM 1:1 or 1:0.5 ratio adding the PCR product. PCR product(s) and ∼8 μCi of [^35^S]-methionine were mixed for a 15-μL PURExpress reaction, followed by incubation at 37 °C for 30 min at 700 rpm shaking. Translation was ceased by the addition of 10 µL of ice-cold 10% trichloroacetic acid (TCA), followed by incubation on ice for at least 30 min. Total protein was sedimented by centrifugation at 4 °C for 5 min at 20,000 g. The supernatant is carefully removed and the pellet was resuspended in a suitable volume of 1× SDS/PAGE sample buffer (134 mM Tris·HCl at pH 8, 13.5% (v/v) glycerol, 3.32% SDS, 0.075% bromophenol blue, 10 mM EDTA and 100 mM DTT) by shaking at 37 °C and 900 rpm for 10 min. The prolyl-tRNA that remains attached due to SecM arrest is digested by the addition of 4 μL of a 4 μg/μL RNase I solution, followed by incubation at 37 °C and 700 rpm for 10 min.

Following a brief centrifugation (2 min at 20,000 g) to remove any remaining insoluble material, the sample is loaded onto an appropriate SDS/PAGE gel (12% Bis-Tris gels were used for large constructs and and 16% Tricine gels were used for small constructs of the library run in MES or Tricine buffer, respectively). Following electrophoresis, the gels are dried onto thick filter paper by heating under vacuum (Bio-Rad model 583 or Hoefer GD 2000), a radioactive molecular weight ladder included in the gel is visualized by spotting the filter paper with a ∼1:1,000 solution of [^35^S]-methionine in 1× SDS/PAGE sample buffer, and the gel is exposed to a phosphorimager screen for 12 h. The screen was imaged using a Fujifilm FLA9000 (50-μm pixels), and densitometry analysis on the resultant raw image (TIFF format) file was carried out using FIJI (ImageJ) software. The densitometry values are quantified using our in-house EasyQuant software and the fraction full-length protein was calculated as mentioned above. See Supplemental Figure S4 C and E, for examples of gels. Independent replicate *in vitro* translation reactions were conducted for all K_V_7.2 CRD constructs.

### Solubility and FRET assay

Protein expression and bacterial lysis was conducted as in FPA. The soluble protein fractions were analyzed in a Fluoromax-3 fluorometer, recording the emission spectra of mTFP1 and mcpVenus fluorescent proteins upon excitation at 458 and 515 nm, respectively. FRET index was calculated as the ratio between the mcpVenus and mTFP1 emission peak amplitudes after exciting at 458 nm.

Protein solubility was analyzed by in-gel densitometry, comparing protein amounts in equal volume of pellet and supernatant fractions. The protein amount in the pellet and in the supernatant was estimated relative to the total protein amount by quantifying the gel bands using FIJI (ImageJ) software. The gels were run as in FPA protocol.

### Single-molecule force spectroscopy assay

Detailed methods for single-molecule sample preparation, optical tweezers assay and data analysis procedure are provided in the Supplementary Materials.

Briefly, variants that showed peaks IV and VII in the *in vivo* FPA (Fig. S1) were adapted for single-molecule assay through the incorporation of a N-terminal amber codon with a flexible linker, to enable the N-terminus to be attached to the DNA handle. SecM sequence was replaced by the stronger SecMstr variant (FSTPVWIWWWPRIRGPP)^43,44^. dsDNA ‘handles’ were prepared by PCR amplification with digoxygenin and biotin 5’-end-modified primers and purified on an agarose gel. The purified handles were incubated with Neutravidin to quench the exposed biotin tag.

Polystyrene beads were sequentially functionalized with dsDNA handles and biotinylated ribosomes by incubating them with the substrate for 20-60 min, followed by washing with TICO buffer (20 mM HEPES-KOH, 10 mM (Ac)_2_Mg, 30 mM AcNH_4_ and 4 mM β-mercaptoethanol at pH 7.4). Expression of the CRD by the bound ribosomes was induced using a ribosome-free version of the PURExpress *in vitro* transcription-translation kit, supplemented with the plasmid of interest and synthetic biotinylated lysine tRNAs. A second sample of beads was functionalized with dsDNA only using the same method.

Single-molecule experiments were performed using a C-Trap. Samples were measured in an environment containing a second ‘measurement’ buffer (10 mM Tris-HCl, 250 mM NaCl and 10 mM CaCl_2_ at pH 7.0), purified CaM (1 µM in the +CaM condition only), and an oxygen radical scavenging system^45^. One RNC-bead and one DNA-bead were trapped. A single-molecule tether was formed, then repeatedly stretched and relaxed until breakage. The resulting force-extension curves were fitted with a twistable worm-like chain model to extract the protein contour length values reported. Specific conformations of the nascent chains were proposed by comparison of the measured contour lengths with theoretical values computed using a structure of the CaM-bound CRD previously reported. All data analysis was performed using custom scripts in Python.

### Statistical analysis

Values are presented as box plot, indicating the median, lower and upper quartiles and minimum and maximum values. The differences between groups were evaluated using the unpaired Student’s t-test or ANOVA with Mann-Whitney post hoc on SigmaStat Statistic (SigmaPlot 11), where values of p < 0.05 were considered significant. The number of replicates in each experiment are indicated in the figures’ legends. In all figures an asterisk, double asterisks, and triple asterisks indicate significance at p < 0.05, p < 0.01, and p < 0.001, respectively.

## Acknowledgements

This research was supported by the Government of the Autonomous Community of the Basque Country (IT1707-22) and the Spanish Ministry of Science and Innovation (PID2021-128286NB-100, PID2020-118814RB-I00), financed by MCIN/AEI/10.13039/501100011033/FEDER, UE, including FEDER funds. A.M-M, S.M-A and E.N. received support from predoctoral (PRE_2018_1_0126, PRE_2021_1_0101) and postdoctoral (POS_2021_1_0017) contracts, respectively, provided by the Basque Government and administered by the University of the Basque Country (UPV/EHU).

## Authors’ contribution

A.V. and A.M-M. conceived the study and participated in its design and coordination. A.M-M., S.M-A., E.N and J.U. carried out experiments and contributed to figure preparation and manuscript preparation. A.M. and G.v.H. contributed to the *in vitro* experiments. J.R.T., V.S., A.K. and S.J.T. performed the force spectroscopy assays. All authors read and approved the final manuscript.

