## Supplemental Text for "Calmodulin acts as a chaperone during co-translational folding of the Kv7.2 channel Calcium Responsive Domain"

### Supplemental Material

#### Supplemental figures

mTurquoise2.1-CRD-SecM-Venus

| Sequence | Residue | |
|---|---|---|
| FALKVQEQHRQKHFEKRRNPAAGLIQSAWRFHVFSTPVWISQHAPIRGSP | 336 | |
| FALKVQEQHRQKHFEKRRNPAAGLIQSAWRFYHVFSTPVWISQHAPIRGSP | 337 | I |
| FALKVQEQHRQKHFEKRRNPAAGLIQSAWRFYAHVFSTPVWISQHAPIRGSP | 338 | |
| FALKVQEQHRQKHFEKRRNPAAGLIQSAWRFYATHVFSTPVWISQHAPIRGSP | 339 | |
| FALKVQEQHRQKHFEKRRNPAAGLIQSAWRFYATNHVFSTPVWISQHAPIRGSP | 340 | II |
| FALKVQEQHRQKHFEKRRNPAAGLIQSAWRFYATNLHVFSTPVWISQHAPIRGSP | 341 | |
| FALKVQEQHRQKHFEKRRNPAAGLIQSAWRFYATNLSHVFSTPVWISQHAPIRGSP | 342 | |
| FALKVQEQHRQKHFEKRRNPAAGLIQSAWRFYATNLSRHVFSTPVWISQHAPIRGSP | 343 | |
| FALKVQEQHRQKHFEKRRNPAAGLIQSAWRFYATNLSRTHVFSTPVWISQHAPIRGSP | 344 | III |
| FALKVQEQHRQKHFEKRRNPAAGLIQSAWRFYATNLSRTDHVFSTPVWISQHAPIRGSP | 345 | |
| FALKVQEQHRQKHFEKRRNPAAGLIQSAWRFYATNLSRTDLHVFSTPVWISQHAPIRGSP | 346 | |
| FALKVQEQHRQKHFEKRRNPAAGLIQSAWRFYATNLSRTDLHSTHVFSTPVWISQHAPIRGSP | 349 | |
| FALKVQEQHRQKHFEKRRNPAAGLIQSAWRFYATNLSRTDLHSTWQYHVFSTPVWISQHAPIRGSP | 352 | |
| FALKVQEQHRQKHFEKRRNPAAGLIQSAWRFYATNLSRTDLHSTWQYYEHVFSTPVWISQHAPIRGSP | 354 | |
| FALKVQEQHRQKHFEKRRNPAAGLIQSAWRFYATNLSRTDLHSTWQYYEFTVHVFSTPVWISQHAPIRGSP | 357 | |
| FALKVQEQHRQKHFEKRRNPAAGLIQSAWRFYATNLSRTDLHSTWQYYEFTVTVPHVFSTPVWISQHAPIRGSP | 360 | |
| FALKVQEQHRQKHFEKRRNPAAGLIQSAWRFYATNLSRTDLHSTWQYYEFTVTVPMYREHVFSTPVWISQHAPIRGSP | 364 | |
| FALKVQEQHRQKHFEKRRNPAAGLIQSAWRFYATNLSRTDLHSTWQYYEFTVTVPMYREDLHVFSTPVWISQHAPIRGSP | 366 | |
| FALKVQEQHRQKHFEKRRNPAAGLIQSAWRFYATNLSRTDLHSTWQYYEFTVTVPMYREDLTPHVFSTPVWISQHAPIRGSP | 368 | |
| FALKVQEQHRQKHFEKRRNPAAGLIQSAWRFYATNLSRTDLHSTWQYYEFTVTVPMYREDLTPGLHVFSTPVWISQHAPIRGSP | 370 | |
| FALKVQEQHRQKHFEKRRNPAAGLIQSAWRFYATNLSRTDLHSTWQYYEFTVTVPMYREDLTPGLKVHVFSTPVWISQHAPIRGSP | 372 | IV |
| FALKVQEQHRQKHFEKRRNPAAGLIQSAWRFYATNLSRTDLHSTWQYYEFTVTVPMYREDLTPGLKVSIHVFSTPVWISQHAPIRGSP | 502 | |
| FALKVQEQHRQKHFEKRRNPAAGLIQSAWRFYATNLSRTDLHSTWQYYEFTVTVPMYREDLTPGLKVSIRHVFSTPVWISQHAPIRGSP | 503 | |
| FALKVQEQHRQKHFEKRRNPAAGLIQSAWRFYATNLSRTDLHSTWQYYEFTVTVPMYREDLTPGLKVSIRAVCVMRFHVFSTPVWISQHAPIRGSP | 510 | |
| FALKVQEQHRQKHFEKRRNPAAGLIQSAWRFYATNLSRTDLHSTWQYYEFTVTVPMYREDLTPGLKVSIRAVCVMRFLVSKRKHVFSTPVWISQHAPIRGSP | 516 | |
| FALKVQEQHRQKHFEKRRNPAAGLIQSAWRFYATNLSRTDLHSTWQYYEFTVTVPMYREDLTPGLKVSIRAVCVMRFLVSKRKFKESHVFSTPVWISQHAPIRGSP | 520 | V |
| FALKVQEQHRQKHFEKRRNPAAGLIQSAWRFYATNLSRTDLHSTWQYYEFTVTVPMYREDLTPGLKVSIRAVCVMRFLVSKRKFKESLRPYDHVFSTPVWISQHAPIRGSP | 525 | |
| FALKVQEQHRQKHFEKRRNPAAGLIQSAWRFYATNLSRTDLHSTWQYYEFTVTVPMYREDLTPGLKVSIRAVCVMRFLVSKRKFKESLRPYDVMDVIHVFSTPVWISQHAPIRGSP | 530 | VI |
| FALKVQEQHRQKHFEKRRNPAAGLIQSAWRFYATNLSRTDLHSTWQYYEFTVTVPMYREDLTPGLKVSIRAVCVMRFLVSKRKFKESLRPYDVMDVIEQYSAHVFSTPVWISQHAPIRGSP | 535 | VII |

hA hTW hB

mTFP1-CRD-L21-SecM-Venus

| Sequence | Residue |
|---|---|
| FALKVQEQHRQKHFEKRRNPAAGLIQSAWRFYATNLSRTDLHSTWQYYEFTVTVPMYREDLTPGLKVSIRAVCVMRFLVSKRKFKESLREFYVFSTPVWISQHAPIRGSP | L21 |

**Figure S1. Amino acid sequences of constructs used for the FPA.** Increasing lengths of the Kv7.2 CRD were cloned upstream of the SecM (*Ec-Ms*) (in green) separated by a two-residues-linker (in gray). In each construct, the amino acid located 30-residues-upstream PTC is highlighted in light blue as a reference. Upstream from this point these are the residues emerging from the ribosome, when the arrest should occur if the nascent chain does not exert force. CRD helices are underlined and colored in blue, green and reed for the hA, hTW and hB, respectively. The number on the right column corresponds to the $K_V$7.2 residue number of the reference amino acids highlighted in light blue. Note that linker connecting hTW and hB, previously described as Δ6L (ΔR373-T501)[4] has been deleted in these constructs.

mTurquoise2.1-CRD-SecM-Venus

| | | |
|---|---|---|
| FALKVQEQHRQKHFEKRRNPAAGLIQSAWRFYATNLSRTDLHSTWQYYEFTVTVPMYREDLTPGLKVHVFSTPVWISQHAPIRGSP | 372 | IV |
| FALKVQEQHRQKHFEKRRNPAAGSGSSGSGSGSTNLSRTDLHSTWQYYEFTVTVPMYREDLTPGLKVHVFSTPVWISQHAPIRGSP | 372 | IV |
| FALKVQEQHRQKHFEKRRNPAAGLIQSAWRFYATNLSRTDLGSGSGSGSGSVTVPMYREDLTPGLKVHVFSTPVWISQHAPIRGSP | 372 | IV |
| FALKVQEQHRQKHFEKRRNPAAGLIQSAWRFYATNLSRTDLHSTWQYYEFTVTVPMYREDLTPGLKVSIRAVCVMRFLVSKRKFKESLRPYDVMDVIEQYSAHVFSTPVWISQHAPIRGSP | 535 | VII |
| FALKVQEQHRQKHFEKRRNPAAGSGSSGSGSGSTNLSRTDLHSTWQYYEFTVTVPMYREDLTPGLKVSIRAVCVMRFLVSKRKFKESLRPYDVMDVIEQYSAHVFSTPVWISQHAPIRGSP | 535 | VII |
| FALKVQEQHRQKHFEKRRNPAAGLIQSAWRFYATNLSRTDLGSGSGSGSGSVTVPMYREDLTPGLKVSIRAVCVMRFLVSKRKFKESLRPYDVMDVIEQYSAHVFSTPVWISQHAPIRGSP | 535 | VII |

**Figure S2. Amino acid sequences of constructions corresponding to peak IV and VII and its GSG-substituted analogs.** The same color code as in Fig S1 has been used and the GSG linker was underlined in dark gray.

mTFP1-CRD-Venus

| | |
|---|---|
| FALKVQEQHRQKHFEKRRNPAAGLIQSAWRFYATNLSRTDLHSTWQYYERTVTVPMYREDLTPGLKVSIRAVCVMRFLVSKRKFKESLRPYDVMDVIEQYSAGHLDRPVD | hA – hTW – hB |
| FALKVQEQHRQKHFEKRRNPAAGSGSSGSGSGSTNLSRTDLHSTWQYYERTVTVPMYREDLTPGLKVSIRAVCVMRFLVSKRKFKESLRPYDVMDVIEQYSAGHLDRPVD | GSG– hTW – hB |
| FALKVQEQHRQKHFEKRRNPAAGLIQSAWRFYATNLSRTDLGSGSGSGSGSVTVPMYREDLTPGLKVSIRAVCVMRFLVSKRKFKESLRPYDVMDVIEQYSAGHLDRPVD | hA – GSG – hB |
| FALKVQEQHRQKHFEKRRNPAAGLIQSAWRFYATNLSRTDLHSTWQYYERTVTVPMYREDLTPGLKGSGSGSGSGSFLVSKRKFKESLRPYDVMDVIEQYSAGHLDRPVD | hA – hTW –GSG |
| FALKVQEQHRQKHFEKRRNPAAGLIQSAWRFYATNLSRTDLHSTWQYYERTVTVP | hA – hTW |

**Figure S3. Amino acid sequences of folding biosensors.** Kv7.2 CRD was flanked between mTFP1 and mcpVenus in the N- and C-terminals, respectively. In independent biosensors, each helix of the CRD was replaced with a GSG linker (in gray). CRD helices are underlined and colored in blue (hA), green (hTW) and red (hB). Note that the linker connecting hTW and hB is deleted in all constructs (ΔR373-T501)[4].

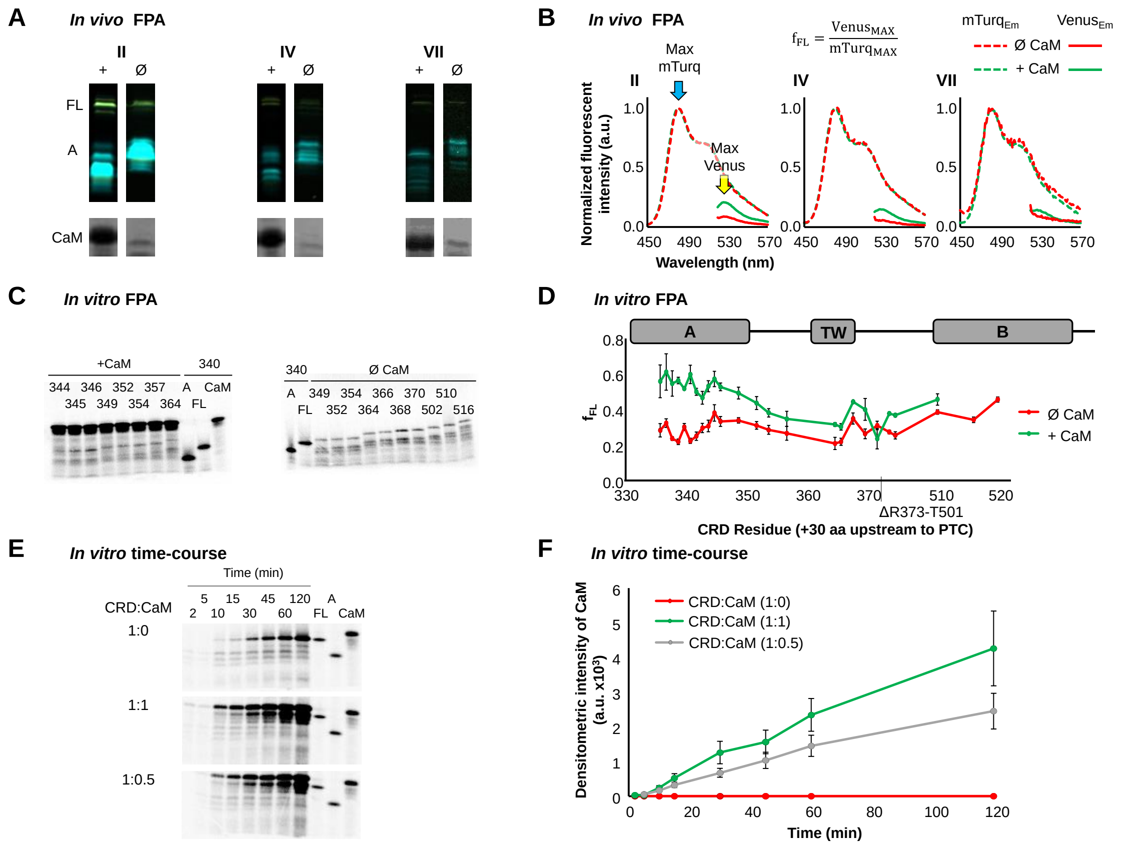


**Figure S4.** ***In vivo* and *in vitro* FPA.** **(A)** Fluorescent image of an SDS-PAGE gel of unboiled bacterial extract for the FPA CRD variants generating peaks II, IV and VII, respectively, alone (Ø) and co-expressed with CaM (+). The Turquoise emission is represented as cyan and the Venus one as yellow. Note that FL fraction expresses both fluorescent proteins and thus the bands of the proteins are colored in green, while the A fraction only expresses Turquoise. CaM expression is indicated at the bottom by Commassie staining. **(B)** Normalized emission spectra of the FPA CRD variants generating peaks II, IV and VII, respectively, alone (Ø, in red) and co-expressed with CaM (in green). Turquoise and Venus emission spectra are indicated by dashed and continuous lines, respectively. The formula for computing $f_{FL}$ is shown on top. **(C)** Radioactive gels of the *in vitro* FPA for the different variants, indicated with the length, with and without co-expressing CaM (left and right, respectively). Full-length (FL), arrested (A) control variant products and CaM expression alone were charged in the gels as controls. **(D)** *In vitro* FPA of CRD variants expressed alone and co-expressed with CaM (red and green, respectively). A schematic representation of the CRD is presented on top. Bars indicate SEM (n = 3). **(E)** Radioactive gels of the *In vitro* time-course pulling-force assay of CRD without and with different amounts of CaM cDNA (0 ng, 100 ng and 50 ng of CaM for top, middle and bottom gels, respectively. Time is indicated on top. **(F)** Fluorimetric $f_{FL}$ for the *in vitro* time-course pulling-force assay of CRD without and with different amounts of CaM cDNA (0 ng, 50 ng and 100 ng, are represented in red, gray and green, respectively.

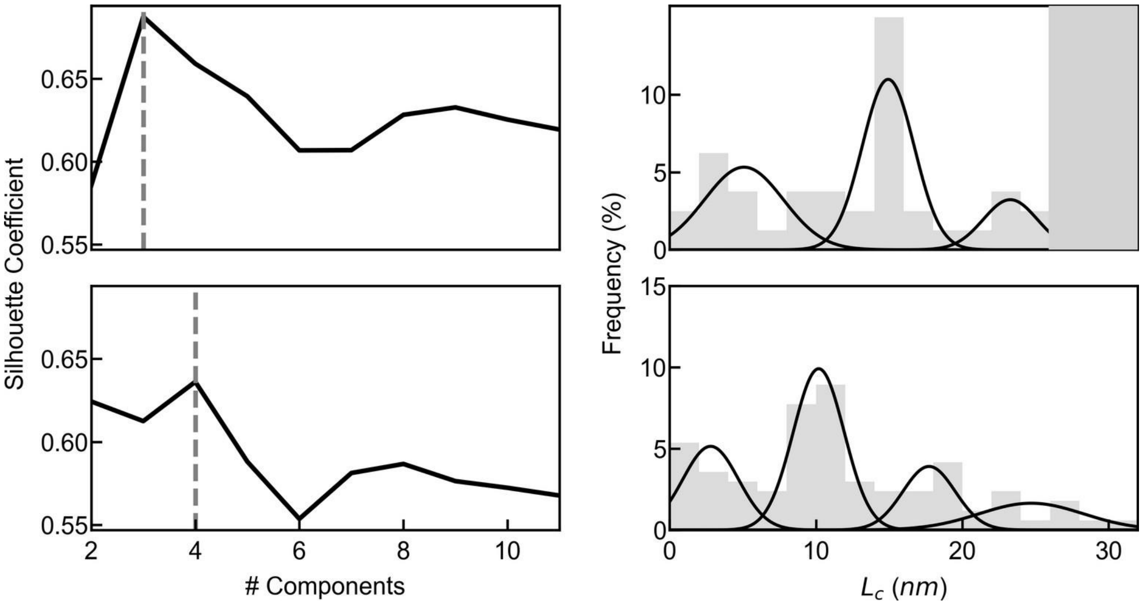


**Figure S5. Experimental Lc values were calculated with a Gaussian mixture fit of the histograms shown in Figure 4.** Each histogram was fitted with a Gaussian Mixture model with 2-11 components, and the model with the highest silhouette coefficient [45] was selected. Left: plots of silhouette coefficient vs. number of components – the optimal number of components is indicated with a gray dashed line. Right: the +CaM histograms for each of the two constructs - the black lines indicate the probability density functions for each component of the best-fit Gaussian Mixture model. The fit components for the $IV_{SM}$ construct are 5.1 ± 2.7 nm, 14.9 ± 1.8 nm, and 23.3 ± 1.8 nm; the components for the $VII_{SM}$ construct are 2.8 ± 1.9 nm, 10.2 ± 1.7 nm, 17.7 ± 1.8 nm, and 24.7 ± 3.6 nm. The dark gray box in the top right plot indicates contour lengths longer than the construct itself, which therefore cannot be reached through mechanical unfolding.

**Table S1. Theoretical and experimental Lc values for each possible hairpin structure.** Columns contain, in order: the state number corresponding with those in Figure 4E; the construct name; each pair of α-helices for each construct which may bind into a hairpin structure; the number of amino acids on the N-terminal side of the hairpin, which are free to fully extended under applied force; similar free amino acids on the C-terminal side of the hairpin outside the ribosome exit tunnel; the theoretical Lc value, calculated using Equation 1; and the experimental Lc value (mean ± standard deviation) obtained from force spectroscopy experiments. Experimental values were calculated with a Gaussian mixture fit of the histograms shown in Figure 4D (Figure S5). For each state which is observed in force spectroscopy experiments, the experimental measurements are in close agreement with theoretical values.

| # | construct | hairpin | N-term (aa) | C-term (aa) | theor. $L_c$(nm) | expt. $L_c$ (nm) |
|---|---|---|---|---|---|---|
| 1 | $IV_{SM}$ | hA / hTW | 23 | 22 | 16.4 | 14.9 ± 1.8 |
| 2 | $VII_{SM}$ | hA / hB | 23 | 0 | 8.9 | 10.2 ± 1.7 |
| 3 | $VII_{SM}$ | hTW / hB | 47 | 0 | 17.1 | 17.7 ± 1.8 |
| - | $VII_{SM}$ | hA / hTW | 23 | 42 | 23.2 | 24.7 ± 3.6 |

#### Supplementary Methods

*Plasmid construction*

Variants that gave peaks IV and VII in the *in vivo* FPA (Fig. S1) were adapted for single-molecule assay through the incorporation of a N-terminal amber codon with a flexible linker, to enable the N-terminus to be attached to the DNA handle. SecM sequence was replaced with the more force-resistant SecMstr (FSTPVWIWWWPRIRGPP)[42,43]. The modified sequences were cloned into pRSET plasmid between the BamHI and XhoI restriction sites.

*Isolation of biotinylated ribosomes*

Ribosomes from Can20/12E37, an RNAse deficient *E. coli* K-12 strain[45], were biotinylated *in vivo* at the uL4 ribosomal protein and isolated as previously described[46]. Ribosomal activity was verified by synthesis of GFP emerald (GFPem) using a ribosome-free *in vitro* transcription-translation system[28] (PURExpress® Δ Ribosome Kit; New England Biolabs, E3313S) supplemented with the isolated ribosomes and a DNA plasmid encoding GFPem. Synthesis of GFPem was confirmed by fluorescence measurement using a QM-7 spectrofluorometer (Photon Technology International).

*Coupling of ribosomes to beads with DNA handles*

Five kbp double-stranded DNA (dsDNA) 'handles' were prepared by PCR amplification with digoxygenin and biotin 5'-end-modified primers and purified on an agarose gel. The resulting PCR fragments were incubated with a 200-fold excess of NeutrAvidin (NTV; Thermo Scientific, 31000) on a rotary mixer for 24 h at 4 °C.

Before each measurement, two batches of bead mixture were prepared with NTV-DNA handles (1.4 fmol), Ø 2.1 µm anti-digoxygenin-coated polystyrene beads (Spherotech, DIGP-20-2; 0.1% w/v; 2 µL) and TICO buffer (20 mM HEPES-KOH, 10 mM $(Ac)_2Mg$, 30 mM $AcNH_4$ and 4 mM β-mercaptoethanol at pH 7.4; 12 µL) and incubated on a rotary mixer at 4 °C. The first batch (DNA-beads) was stored on a rotary mixer at 4 °C. Immediately before measurement, the DNA-bead mixture was further diluted in TICO buffer (288 µL). The second batch was pelleted and re-suspended in TICO (50 µL); pelleted again and re-suspended in TICO (20 µL) supplemented with biotinylated ribosomes (30 pmol) and RNase Inhibitor Murine (New England Biolabs, M0314S; 10 units). This mixture was incubated on a rotary mixer for 45 min at 4 °C. The resulting ribosome-bead mixture was pelleted and re-suspended in TICO (50 µL) to remove excess ribosomes.

*Generation of stalled ribosome-nascent chain complexes (RNCs)*

A ribosome-free *in vitro* transcription-translation system[28] (PURExpress® Δ Ribosome Kit; New England Biolabs, E3313S) was prepared with a final volume 12 µL. This PURE mixture was supplemented with synthetic tRNAs encoding the UAG stop codon, pre-charged with biotinylated lysine amino acids (Hölzel, PRX-CLD04; 120 pmol) and the circular plasmid encoding the construct of interest (60 fmol). The ribosome-bead mixture was pelleted, resuspended in the modified PURE mixture, spun down on a microfuge for 3 s and incubated for 20 min at 37 °C to generate bead-bound, stalled ribosome-nascent chain complexes (RNC-beads). Immediately before measurement, the DNA-bead mixture was diluted further in TICO (288 µL).

*Preparation of measurement buffer*

Single-molecule experiments were performed in an environment containing an oxygen radical scavenging system[44] (3 units mL-1 pyranose oxidase, Merck, P4234; 90 units mL-1 catalase, Merck, C9322; 50 mM glucose) and purified CaM (1 µM in the +CaM condition only) in a buffer containing 10 mM Tris-HCl, 250 mM NaCl and 10 mM $CaCl_2$ at pH 7.0, as previously reported[47,48].

This 'measurement buffer' was prepared, pelleted and the supernatant collected immediately before the measurement.

*Optical tweezers setup & force spectroscopy experiments*

The single-molecule experiments were performed using a C-Trap (Lumicks). In brief, the instrument consists of two optical traps formed by a single high-intensity, polarisationstable 1064 nm laser split into two orthogonally-polarised beams. Samples are manipulated in a monolithic laminar flow cell with five separate flow channels controlled by a passive pressure-driven microfluidic system. The microfluidic system is modified with a custom cooling setup enabling samples to be stored at 4 °C before injection into the flow cell, to improve sample lifetime.

During the experiment, a RNC-bead was collected from one laminar flow channel in one trap, and a DNA-bead from a second channel in the other. The beads were moved to a third channel containing the measurement buffer prior to tether formation. The beads were repeatedly brought into close proximity and back until a slight increase in measured force upon retraction indicated formation of a tether. Single tethers were identified according to three criteria: the tether length (ca. 3.4 µm corresponding to the two 5 kbp dsDNA handles); presence of the characteristic twist-stretch bending motif above 35 pN[49]; and presence of an unfolding transition with length matching the expected length of the nascent CRD (28 nm and 34 nm for the hA-hTW and hA-hTW-hB constructs, respectively). Measurements were taken in a cycling 'force spectroscopy' mode, where the steerable trap was moved at a constant rate of 0.1 $\mu m \cdot s^{-1}$ between a minimum trap separation of 2.2 µm and a maximum applied force of 35 – 65 pN repeatedly until tether breakage.

*Data analysis*

Force-distance data were collected at 50 kHz and decimated to 500 Hz prior to analysis. For each bead pair, the optical traps were calibrated by fitting a Lorentzian function to the power spectrum of the Brownian motion of the trapped beads12, The trapping laser intensity was kept constant for all measurements, resulting in trap stiffness values of 335 ± 108 pN µm-1. Each force-extension curve was identified and fit with two worm-like chain (WLC) models in series: a twistable WLC11 for the DNA component, and the Odijk approximation for an inextensible WLC13 for the protein component. For these fits, the DNA contour length ($L_C$), protein persistence length (Lp) and twist-stretch coupling critical force were held constant at 3.4 µm, 0.75 nm and 30.6 pN respectively. The DNA Lp, DNA stretch modulus (St), DNA twist rigidity (C), and the twist-stretch coupling parameters g0 and g1 were fit, yielding average values of 33.5 ± 10.8 nm, 1200 ± 460 pN nm-1, 460 ± 87 pN nm2, -295 ± 101 pN nm, and 11.0 ± 2.8 nm respectively. Folding events were identified and quantified using a semi-automated K-means clustering algorithm. All calibration, fitting, and folding event identification was performed using custom scripts in Python.

*Determination of theoretical contour lengths*

The protein contour length values measured upon mechanical unfolding of the nascent CRD were interpreted by comparison to the solution structure of the CaM-bound /CRD complex previously reported (PDB 6FEG)[4]. The CRD structure formed upon binding of CaM comprises a central hairpin region flanked by unstructured polypeptide on both the N- and C-termini. Therefore, the contour length ($L_c$) is given by Equation 1:

$$L_c = L_{N-term} + x_{hairpin} + L_{C-term} \quad \text{(Equation 1)}$$

where $L_{N\text{-}term}$ and $L_{C\text{-}term}$ are the contour lengths of the N- and C-terminal unstructured regions respectively, and $x_{hairpin}$ is the native extension of the hairpin region. For all proposed contour length states, $x_{hairpin}$ was taken as the Euclidean distance between the Cα atoms of the His35 and Arg113 residues in the crystal structure, 1.1 nm. Taking as an example the aforementioned crystal structure, where the CaM-bound hairpin is formed between the hA and hB α-helices leaving 2 aa and 34 aa unstructured regions at the N- and C-termini respectively, the theoretical protein contour length is:

$$Lc = (2\ aa \cdot 0.34\ nm \cdot aa^{-1}) + 1.1\ nm + (34\ aa \cdot 0.34\ nm \cdot aa^{-1}) = 13.4\ nm$$

#### Supplemental text 1

The library of clones used for the *in vivo* FPA were cloned in pET19b plasmid using Gibson Assembly for *in vitro* transcription/translation using PURExpress commercial system. As the N- and C-terminal of the designed constructs were identical, this technique allowed carrying out a high throughput cloning using the same four primers for all the inserts: two for the vector and two for the insert. The forward and reverse primers of the insert were designed with an overhang of 25 nucleotides hybridizing in the vector upstream and downstream the cloning site, respectively. In addition, both primers were designed with an insert-hybridizing sequence for the 5'-end of the CRD and 3'-end of SecM coding sequences, for forward and reverse primers, respectively with lengths adjusted to Tm ≥ 60 °C. As a vector, a previously employed clone for *in vitro* transcription/translation assay was used, which consisted on a 3'-end gene sequence encoding 23 residues (a LepB P2 domain-derived sequence, GSSDKQEGEWPTGLRLSRIGGIH) C-terminal. The reverse primer of the vector was designed upstream the cloning site adjusting the length to Tm ≥ 60 °C and without overhang. For radiolabeling detection after *in vitro* expression for FPA it is important to eliminate the methionines after SecM sequence, so that both full-length and arrested proteins present the same radioactive intensities. Thus, Venus needed to be substituted. For that, the forward primer of the vector was designed with an overhand of 25 nucleotides hybridizing with the 3'-end of the insert (SecM), and an additional sequence hybridizing with the leptin-coding-zone of the vector adjusting the length to Tm ≥ 60 °C (**Figure S6**). Therefore, this library of constructs cloned in pET19b consisted of increasing lengths of CRD cloned upstream SecM-LepB coding sequence.

The vector and all the insert clones were amplified by PCR using Q5® high-fidelity polymerase (**Table S2**). After checking that the products were correctly amplified using for that 5 µL of PRC product to run an agarose gel, the rest of the product was digested with DnpI to eliminate the templates in vector and inserts clones for 1 h at 37 °C. Next, DnpI enzyme was inactivated at 80 °C for 20 minutes. Gibson reaction was then carried out mixing 3 and 2 µL of insert and vector PCR products, respectively, and 5 µL of Gibson reaction mix (**Table S3**) for 1 h at 50 °C. Finally, the Gibson reaction products were transformed in *E. coli* DH5α strain, single colonies were inoculated in liquid medium and DNA miniprecipitations were carried out. New clones were sequenced in Eurofins Genomics as quality control.

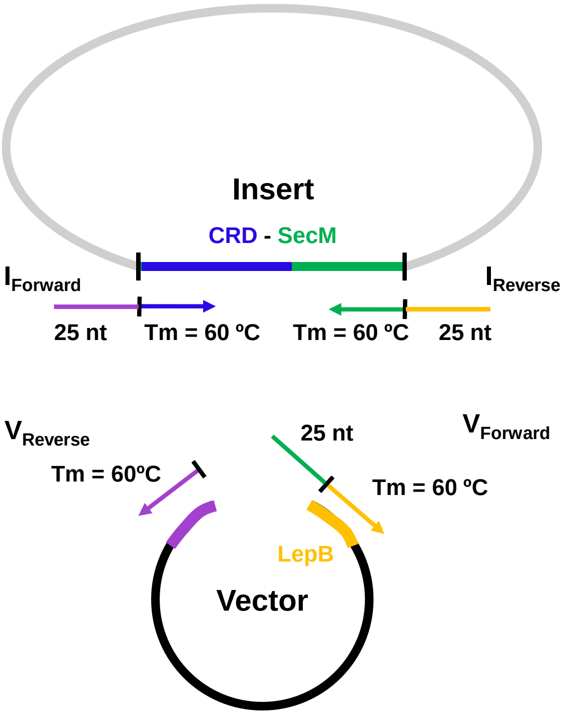


**Figure S6. Primer design for Gibson assembly.** Insert and vector primers are indicated with I and V, respectively, and nt refers to nucleotides. Follow the color code for primer-insert or primer-vector hybridizing sequences, for example, the yellow parts of $I_{Reverse}$ and $V_{Forward}$ hybridize with LepB-coding-sequence of the vector shown also in yellow.

**Table S2. PCR conditions.** Top, reaction conditions. Bottom, PCR protocols for vector and inserts clones amplification.

| DNA (1 µg/µL) | Forward (10 µM) | Reverse (10 µM) | Q5 Buffer | dNTPs (10 mM) | Q5 | $H_2O$ | $V_f$ = 15 µL |
|---|---|---|---|---|---|---|---|
| 0.3 | 0.8 | 0.8 | 4 | 0.4 | 0.1 | 13.6 | (µL) |

| PCR conditions | | | | | |
|---|---|---|---|---|---|
| Vector | | | Insert | | |
| 1. | 98 °C | 1:00 | 1. | 98 °C | 1:00 |
| 2. | 98 °C | 0:20 | 2. | 98 °C | 0:20 |
| 3. | 60 °C | 0:10 | 3. | 60 °C | 0:10 |
| 4. | 72 °C | 4:00 | 4. | 72 °C | 2:00 |
| 5. | GOTO 2 | X20 | 5. | GOTO 2 | X20 |
| 6. | 4 °C | ∞ | 6. | 4 °C | ∞ |

**Table S3. Gibson reaction mix.** Top, at-home made Gibson reaction mix protocol. Bottom, 5X ISO buffer preparation protocol.

| Gibson Raction Mix | | |
|---|---|---|
| **Component** | **Volume** | **Reference** |
| 5X ISO Buffer | 320 µL | See below |
| T5 Exonuclease | 0.64 µL of 10 U/µL (6.4 U) | NEB M0363S |
| Phusion polymerase | 20 µL of 2 U/µL (40 U) | NEB M0530S |
| Taq ligase | 160 µL of 40 U/µL (6400 U) | NEB M0208L |
| Milli-Q $H_2O$ | Up to 1.2 mL final volume | |

| ISO Buffer (5X) | | | |
|---|---|---|---|
| **Component** | **[Final]** | **Volume** | **Reference** |
| Tris HCl, pH 7.5 | 0.5 M | 3 mL of 1 M stock | Sigma T1503 |
| $MgCl_2$ | 50 mM | 60.99 mg | Sigma M0250 |
| DTT | 50 mM | 46.275 mg | Sigma D9779 |
| PEG-8000 | 250 g / L | 1.5 g | Sigma P5413 |
| $NAD^+$ | 5 mM | 19.9 mg | Sigma N7003 |
| dGTP | 10 mM | 60 µl of 100 mM | Thermo R0181 |
| dATP | 10 mM | 60 µl of 100 mM | Thermo R0181 |
| dTTP | 10 mM | 60 µl of 100 mM | Thermo R0181 |
| dCTP | 10 mM | 60 µl of 100 mM | Thermo R0181 |
| Milli-Q $H_2O$ | | To 6 mL final | |
